# APOA1 links a late fetal epithelial program to colonic maturation and injury responses

**DOI:** 10.64898/2026.09.24.754040

**Authors:** Aline Böniger, Valerie Georgette Katimi Varela, Dillon King, Tomas Valenta, Giulia Moro, Costanza Borrelli, Michael David Brügger, Patrick Diener, Andreas E. Moor, Hassan Fazilaty

## Abstract

Developmental programs establish tissue identity and organization, but how their reactivation contributes to adult tissue responses remains poorly understood. Here, we identify a functional contribution of apolipoprotein A-I (APOA1), which marks a late fetal colonic epithelial state that reappears after injury. Spatial and temporal profiling localized *Apoa1* transcription predominantly to epithelial cells at the tips of developing proximal colonic folds. *Apoa1* loss delayed fold maturation and altered the abundance and composition of developing enteroendocrine populations, linking this apolipoprotein to mucosal architecture and epithelial differentiation. APOA1 protein closely interacted with WNT5A during development and injury, while *Apoa1* deficiency increased *Wnt5a* expression and altered its epithelial protein distribution. Following colonic injury, single-cell and spatial profiling identified genotype-associated transcriptional changes involving stress responses, barrier and immune functions, and metabolism, with a shared component across males and females. These changes accompanied reduced enteroendocrine representation and increased representation of inflammation-associated macrophages. Finally, regional and compartmental expression analyses, together with hindgut explant perturbation, identified HOXB7 transcription factor as a regulator of developmental *Apoa1* expression. Together, these findings establish functional relevance for a component of a reactivated late fetal epithelial program, connecting APOA1 to colonic maturation and the epithelial and inflammatory organization of injured tissue.

## Introduction

Intestinal development and repair require coordinated changes in epithelial cell identity and tissue organization. During development, these processes establish the specialized lineages and architecture of the mature mucosa. Following injury, epithelial cells can adopt transient regenerative states that share features with fetal intestinal epithelium^1–6^. This recurrence raises a question: what do genes shared by developmental and regenerative states contribute to tissue maturation and repair?

We previously compared the cellular and transcriptional landscapes of developing mouse colon with those of healthy and injured adult colon and identified embryonic expression programs that reappeared after damage^7^. One epithelial population, annotated as embryonic enterocyte progenitors, emerged during late fetal development, was not identified in healthy adult, and reappeared after injury. Expression of apolipoprotein A-I (APOA1) marked this population in both settings. This pattern prompted us to ask whether APOA1 contributes to the tissue processes associated with its appearance.

APOA1 is the principal protein component of high-density lipoprotein (HDL). It is produced mainly by the liver and small intestine and participates in lipid transport^8^; constitutive *Apoa1* deletion markedly reduces circulating HDL cholesterol^9^. Earlier studies established that human fetal colon synthesizes APOA1 and that *Apoa1*-deficient mice develop more severe experimental colitis^10–12^. Its role in colonic development, however, remains unknown, as do the cellular changes associated with its loss during injury.

Here, we define the spatial and temporal pattern of APOA1 expression and examine the consequences of constitutive *Apoa1* deficiency during colonic development and injury. We identify changes in mucosal architecture and enteroendocrine composition, characterize epithelial and immune features of the injured colon using single-cell and spatial transcriptomics, and investigate the relationship between APOA1 and WNT5A. Finally, embryonic tissue perturbation and reporter assays support HOXB7 as a candidate regulator of *Apoa1* expression. These findings connect a protein expressed by a recurring developmental epithelial state to colonic maturation and the response to injury.

## Results

### APOA1 expression defines a transient epithelial domain in developing proximal colonic folds

To define the anatomical distribution of the APOA1-associated epithelial state, we examined the mouse hindgut across late embryonic and early postnatal development. APOA1 immunoreactivity was weak or undetectable at E16.5, became detectable at E17.5, and was prominent in the proximal hindgut at E18.5. Within the epithelium, APOA1 was concentrated toward the luminal tips of developing mucosal folds. Epithelial staining declined after birth and was weak or undetectable by P5 (Fig. 1A). Comparison of proximal and distal E18.5 hindgut sections confirmed the strong regional enrichment of APOA1 protein in the proximal epithelium (Fig. 1B). Three-dimensional whole-mount imaging further resolved the organization of the proximal mucosal surface and localized APOA1-positive epithelial cells along the upper regions of the folds (Fig. 1C).

**Figure 1.**
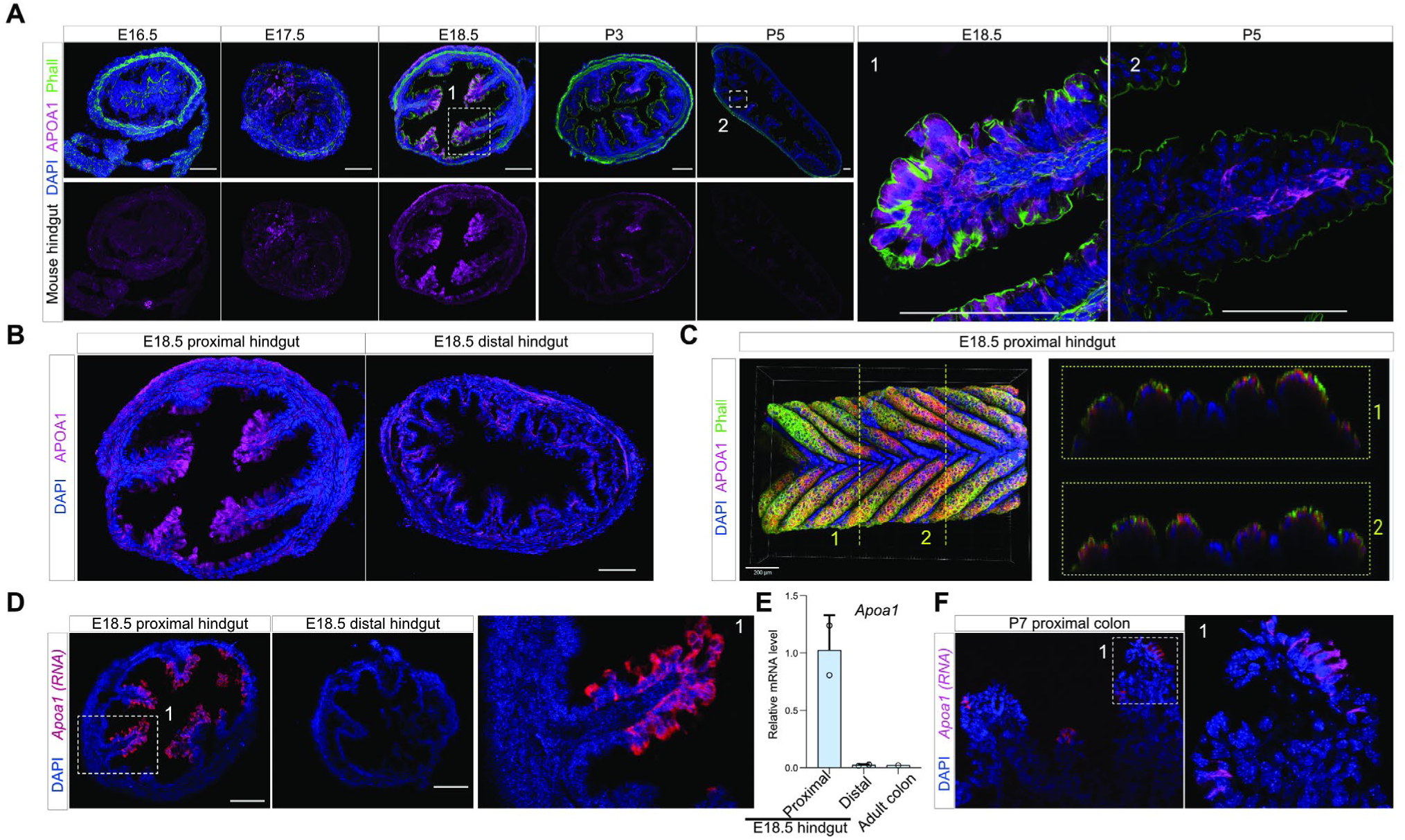
Developmental distribution of APOA1 in the mouse hindgut and colon. (A) Immunofluorescence staining of mouse hindgut sections at E16.5, E17.5, E18.5, P3, and P5. Merged images show APOA1 (magenta), phalloidin (green), and DAPI (blue); the lower row shows the APOA1 channel. Enlarged views of the boxed regions at E18.5 and P5 are shown on the right. (B) APOA1 immunofluorescence staining of proximal and distal E18.5 hindgut sections. (C) Three-dimensional reconstruction of an E18.5 proximal hindgut whole mount stained for APOA1 (magenta), phalloidin (green), and DAPI (blue). Orthogonal sections through the indicated positions 1 and 2 are shown on the right. (D) RNAscope detection of *Apoa1* transcripts (magenta) in proximal and distal E18.5 hindgut sections. The boxed region in the proximal hindgut is shown at higher magnification. (E) Relative *Apoa1* mRNA levels measured by quantitative RT–PCR in E18.5 proximal hindgut, E18.5 distal hindgut, and adult colon. Individual data points are shown. Scale bars are indicated in the images. (F) RNAscope detection of *Apoa1* transcripts in the P7 proximal colon, with an enlarged view of the boxed region. Nuclei were counterstained with DAPI.

We next used RNA in situ hybridization to identify the local source of APOA1 and compare it with the protein distribution. At E18.5, *Apoa1* transcripts were detected predominantly in proximal epithelial cells, particularly near fold tips, with little or no signal in the examined distal sections (Fig. 1D). Quantitative RT–PCR confirmed substantially higher *Apoa1* mRNA levels in E18.5 proximal hindgut than in distal hindgut or adult colon (Fig. 1E). Whereas *Apoa1* transcription was concentrated in the epithelium, APOA1 protein extended into the underlying tissue (Fig. 1A–D), consistent with its known secreted nature. Despite the postnatal decline in epithelial protein staining, *Apoa1* transcripts remained detectable in proximal epithelial cells at P7 (Fig. 1F). Together, these observations identify the proximal epithelium as a principal site of local *Apoa1* expression and show that the distributions of its RNA and protein diverge during maturation.

#### *Apoa1* deficiency alters colonic fold maturation and enteroendocrine composition

The transient expression of APOA1 at developing fold tips prompted us to examine its contribution to colonic maturation using an *Apoa1* knockout (KO) model^9^. At E18.5, three-dimensional whole-mount reconstructions and tissue sections revealed shorter proximal mucosal folds in KO mice than in wild-type (WT) controls (Fig. 2A). APOA1 immunoreactivity was undetectable in knockout tissue, supporting the specificity of the staining. Quantification across development showed reduced proximal fold length at the examined stages from E17.5 through P3, whereas no significant difference was detected at P10 (Fig. 2B). This developmental trajectory is consistent with a transient delay in proximal fold maturation.

**Figure 2.**
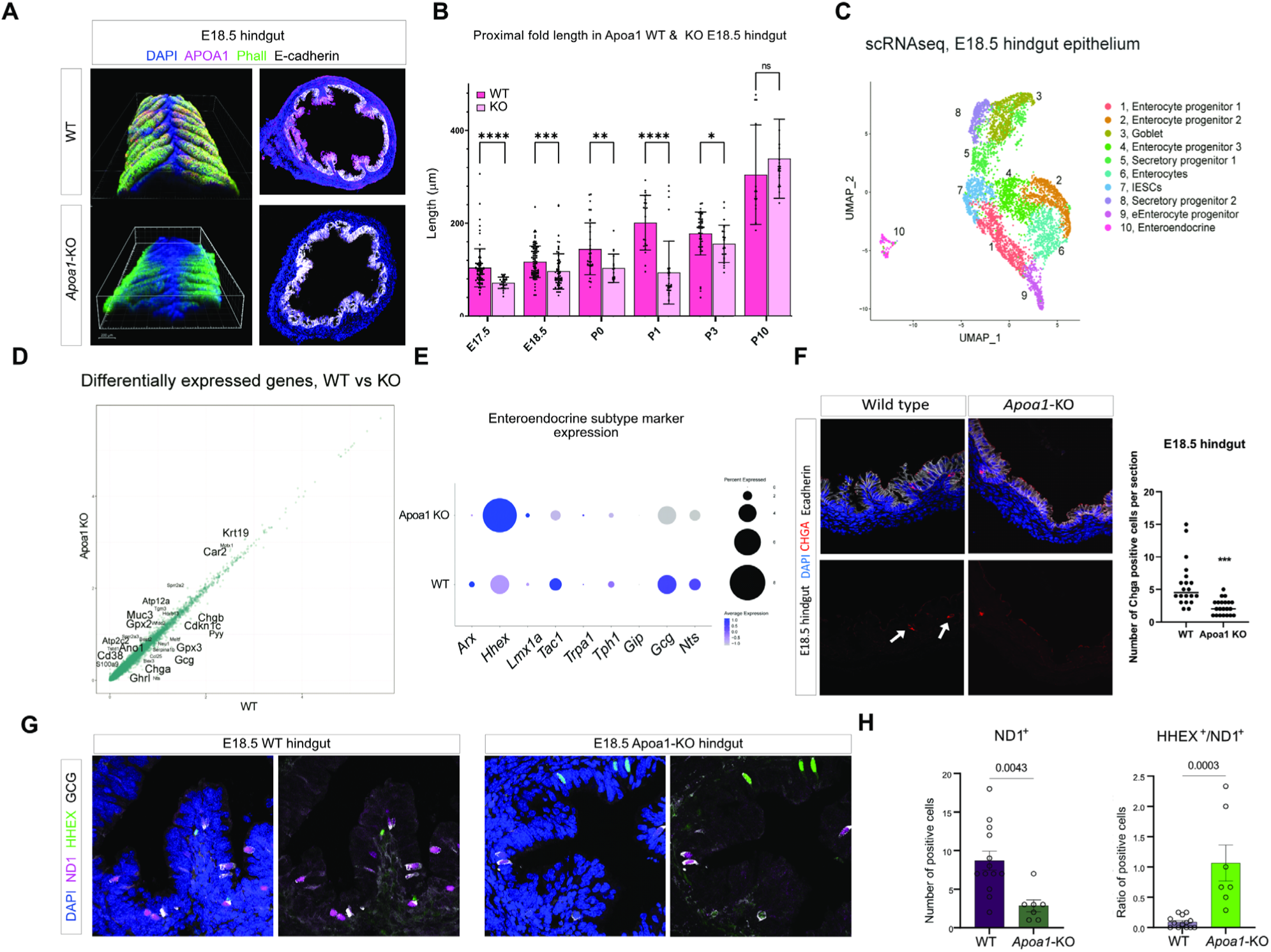
Morphological and enteroendocrine characterization of Apoa1-deficient developing hindgut. (A) Three-dimensional reconstructions and corresponding tissue sections of E18.5 proximal hindgut from wild-type and *Apoa1*-KO mice. Samples were stained for APOA1 (magenta), phalloidin (green), E-cadherin (white), and DAPI (blue). (B) Quantification of proximal mucosal fold length in wild-type and *Apoa1*-KO hindgut or colon at E17.5, E18.5, P0, P1, P3, and P10. Individual measurements and summary statistics are shown. (C) UMAP representation of epithelial cells from E18.5 wild-type and *Apoa1*-KO hindgut analyzed by single-cell RNA sequencing. Cells are colored according to the indicated epithelial cell-state annotations. (D) Comparison of mean gene expression between wild-type and *Apoa1*-KO epithelial cells. Selected differentially expressed genes are labeled. (E) Dot plot showing the expression of selected enteroendocrine subtype markers in wild-type and *Apoa1*-KO epithelial cells. Dot size represents the percentage of expressing cells, and color represents average scaled expression. (F) CHGA immunofluorescence staining of E18.5 wild-type and *Apoa1*-KO hindgut sections. Merged images show CHGA (red), E-cadherin (white), and DAPI (blue); the lower images show the CHGA channel. Arrows indicate CHGA-positive cells. The graph shows the number of CHGA-positive cells per section. (G) Representative E18.5 wild-type and *Apoa1*-KO hindgut sections stained for ND1 (magenta), HHEX (green), GCG (white), and DAPI (blue). Individual and merged fluorescence channels are shown. (H) Quantification of ND1-positive cells and the ratio of HHEX-positive to ND1-positive cells in E18.5 wild-type and *Apoa1*-KO hindgut. Individual data points and summary statistics are shown. Statistical significance is indicated in the corresponding graphs.

To determine which epithelial features accompanied this architectural phenotype, we analyzed single-cell transcriptomes from WT and *Apoa1*-KO E18.5 hindgut epithelium. Clustering resolved the major epithelial progenitor and differentiated populations, including embryonic enterocyte progenitors and enteroendocrine cells (Fig. 2C). Comparison of gene expression highlighted differences in enteroendocrine-associated transcripts, including *Chga*, *Chgb*, and *Gcg* (Fig. 2D). Examination of enteroendocrine subtype markers further identified increased *Hhex* expression in the *Apoa1*-KO cells (Fig. 2E). HHEX is a transcription factor associated with somatostatin-producing intestinal enteroendocrine D cells, suggesting that the endocrine changes might involve differences in composition as well as abundance^13^.

We therefore examined the enteroendocrine compartment directly in tissue. Immunostaining for chromogranin A (CHGA), a broadly expressed enteroendocrine marker, revealed fewer CHGA-positive cells per section in E18.5 *Apoa1*-knockout hindgut than in wild-type controls (Fig. 2F). To further assess endocrine composition, we co-immunostained for NEUROD1 (ND1), a transcription factor involved in enteroendocrine differentiation, HHEX, and GCG, the proglucagon precursor associated with L cells (Fig. 2G). KO tissue contained fewer ND1-positive cells, but a greater proportion of these cells expressed HHEX (Fig. 2H). Thus, *Apoa1* deficiency was associated with a generally smaller endocrine compartment and an increased representation of HHEX-positive cells within it. Together, these findings support a contribution of APOA1 to colonic maturation, linking its loss to delayed fold growth and altered enteroendocrine abundance and composition.

#### Apoa1 deficiency alters WNT5A expression and distribution

To investigate molecular changes associated with the developmental phenotype, we measured selected transcripts related to Wnt signaling, lipid metabolism, and lipoprotein receptors in E18.5 hindguts from WTand *Apoa1*-KO mice. This targeted screen identified increased *Lef1* expression in *Apoa1*-deficient hindgut (Fig. 3A). Immunostaining revealed prominent nuclear LEF1 in KO epithelial cells, whereas nuclear epithelial staining was not detected in WT controls, indicating ectopic epithelial expression (Fig. 3B). This observation is consistent with the reported absence of detectable LEF1 from the developing mouse intestinal epithelium under normal conditions^14^. Given the role of LEF1 in Wnt-regulated transcription, we next examined whether its ectopic expression was accompanied by changes in Wnt-ligand expression. Among the ligands analyzed, *Wnt5a* showed the most pronounced increase in KO hindgut (Fig. 3C). Higher relative *Wnt5a* expression was also detected in the colon of adult *Apoa1*-KO mice compared with WT controls, indicating that the expression difference was not confined to embryonic development (Fig. 3D). WNT5A can modulate β-catenin–TCF signaling in a receptor-dependent manner, and LEF1-regulated transcription has been functionally linked to WNT5A/ROR2 signaling during embryonic morphogenesis^15,16^. These observations motivated further investigation of WNT5A abundance, localization, and epithelial responses in the context of *Apoa1* deficiency.

**Figure 3.**
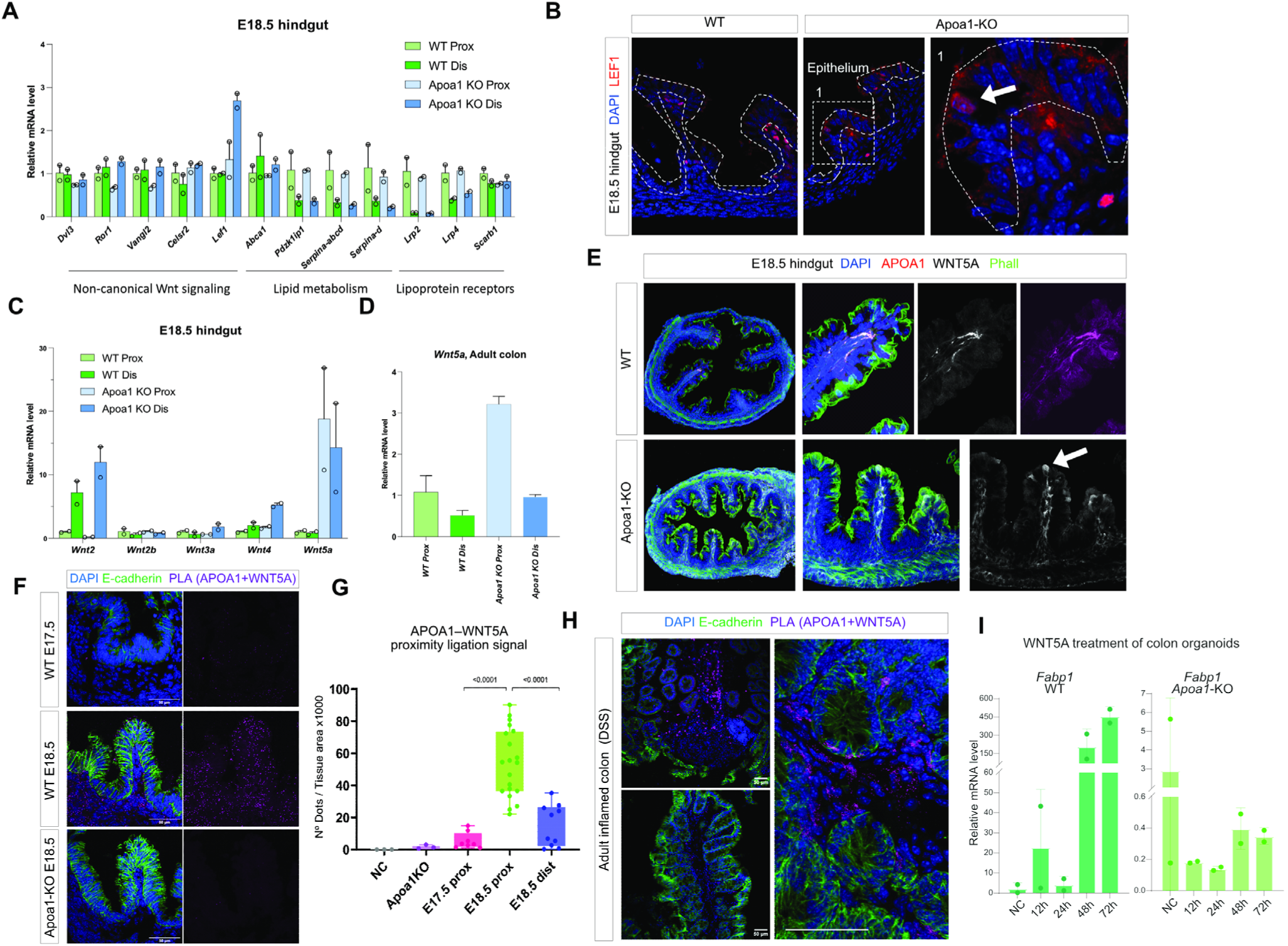
Wnt-related gene expression and APOA1–WNT5A proximity in developing and adult colon. (A) Quantitative RT–PCR analysis of selected genes associated with Wnt signaling, lipid metabolism, and lipoprotein receptors in proximal and distal E18.5 hindgut from wild-type (WT) and *Apoa1*-knockout (KO) mice. (B) LEF1 immunofluorescence staining (red) in WT and *Apoa1*- KO E18.5 hindgut sections, with DAPI counterstaining (blue). Dashed outlines delineate the epithelium. The boxed region is shown at higher magnification on the right; the arrow indicates epithelial LEF1 staining. (C) Quantitative RT–PCR analysis of *Wnt2*, *Wnt2b*, *Wnt3a*, *Wnt4*, and *Wnt5a* in proximal and distal E18.5 hindgut from WT and *Apoa1*-KO mice. (D) Relative *Wnt5a* mRNA levels in proximal and distal colon from healthy, untreated adult WT and *Apoa1*-KO mice. (E) Immunofluorescence staining of E18.5 WT and *Apoa1*-KO hindgut for APOA1 (magenta) and WNT5A (white), with phalloidin (green) and DAPI (blue). Overview images, enlarged views, and separate WNT5A and APOA1 channels are shown. (F) Proximity ligation assay (PLA) for APOA1 and WNT5A in WT E17.5, WT E18.5, and *Apoa1*-KO E18.5 hindgut sections. Merged images show PLA signal (magenta), E-cadherin (green), and DAPI (blue); the corresponding PLA channel is shown alongside each image. (G) Quantification of PLA signal normalized to tissue area in the negative control (NC), *Apoa1*-KO tissue, WT E17.5 proximal hindgut, and WT E18.5 proximal and distal hindgut. (H) APOA1–WNT5A PLA in DSS-treated adult colon, with E-cadherin (green) and DAPI (blue). Overview images and corresponding enlarged views are shown. Prox, proximal; Dis, distal. (I) RT-qPCR analysis of *Fabp1* expression in adult colon-derived wild-type and *Apoa1*-knockout organoids following WNT5A treatment for 12, 24, 48, or 72 h, alongside the negative control (NC).

We next examined the spatial relationship between APOA1 and WNT5A. Immunofluorescence showed overlapping protein distributions in WT E18.5 proximal hindgut, whereas KO tissue displayed more prominent epithelial WNT5A staining (Fig. 3E). Together with the increased transcript levels, this observation suggested that *Apoa1* deficiency affects WNT5A expression and its tissue distribution.

To assess whether APOA1 and WNT5A occur physically interact, we performed an in situ proximity ligation assay (PLA). Signals were detected in WT developing hindgut and were largely absent in KO tissue (Fig. 3F). Quantification showed the highest signal in E18.5 proximal hindgut, with lower levels at E17.5 and in E18.5 distal hindgut (Fig. 3G). PLA signals were also detected in damaged adult colon, extending the observed APOA1– WNT5A proximity to the injury setting (Fig. 3H).

To test whether loss of APOA1 alters epithelial responses to WNT5A, we generated organoids from adult WT and *Apoa1*-KO colons and treated them with recombinant WNT5A protein. Previous studies implicate WNT5A in the regulation of colonic epithelial proliferation and tissue organization and suggest a contribution to enterocyte differentiation^17,18^. We therefore assessed *Fabp1*, a marker of the absorptive epithelial lineage used in mouse colon and colonic organoids^19^, as a differentiation-associated readout. During WNT5A treatment, *Fabp1* expression increased markedly at later time points in WT organoids, whereas KO organoids showed no comparable increase (Fig. 3I). These findings suggest divergent regulation of absorptive differentiation in WT and *Apoa1*-deficient epithelium under WNT5A exposure.

Together, these findings link *Apoa1* deficiency to altered WNT5A abundance and tissue distribution and demonstrate APOA1–WNT5A proximity during development and injury. In colonic organoids, *Apoa1* loss was associated with a distinct differentiation-associated expression response under WNT5A treatment. These complementary observations connect APOA1 to both the tissue distribution of WNT5A and epithelial behavior in its presence.

#### *Apoa1* deficiency alters injury-associated transcriptional responses and epithelial–immune organization

The re-emergence of the APOA1-associated epithelial program following colonic injury prompted us to examine the consequences of Apoa1 deficiency in the adult injured colon. We performed single-cell RNA sequencing of colonic tissue from untreated and DSS-treated WT and Apoa1-KO mice, with three biological replicates for each sex, genotype and treatment combination (Fig. 4A). Integration of the 24 individually barcoded samples resolved epithelial, immune and mesenchymal populations, including multiple epithelial states, enteroendocrine cells, two macrophage populations, neutrophils and lymphoid populations (Fig. 4B). The major cellular compartments were recovered in both genotypes, providing a framework for investigating genotype-associated transcriptional and compositional differences.

**Figure 4.**
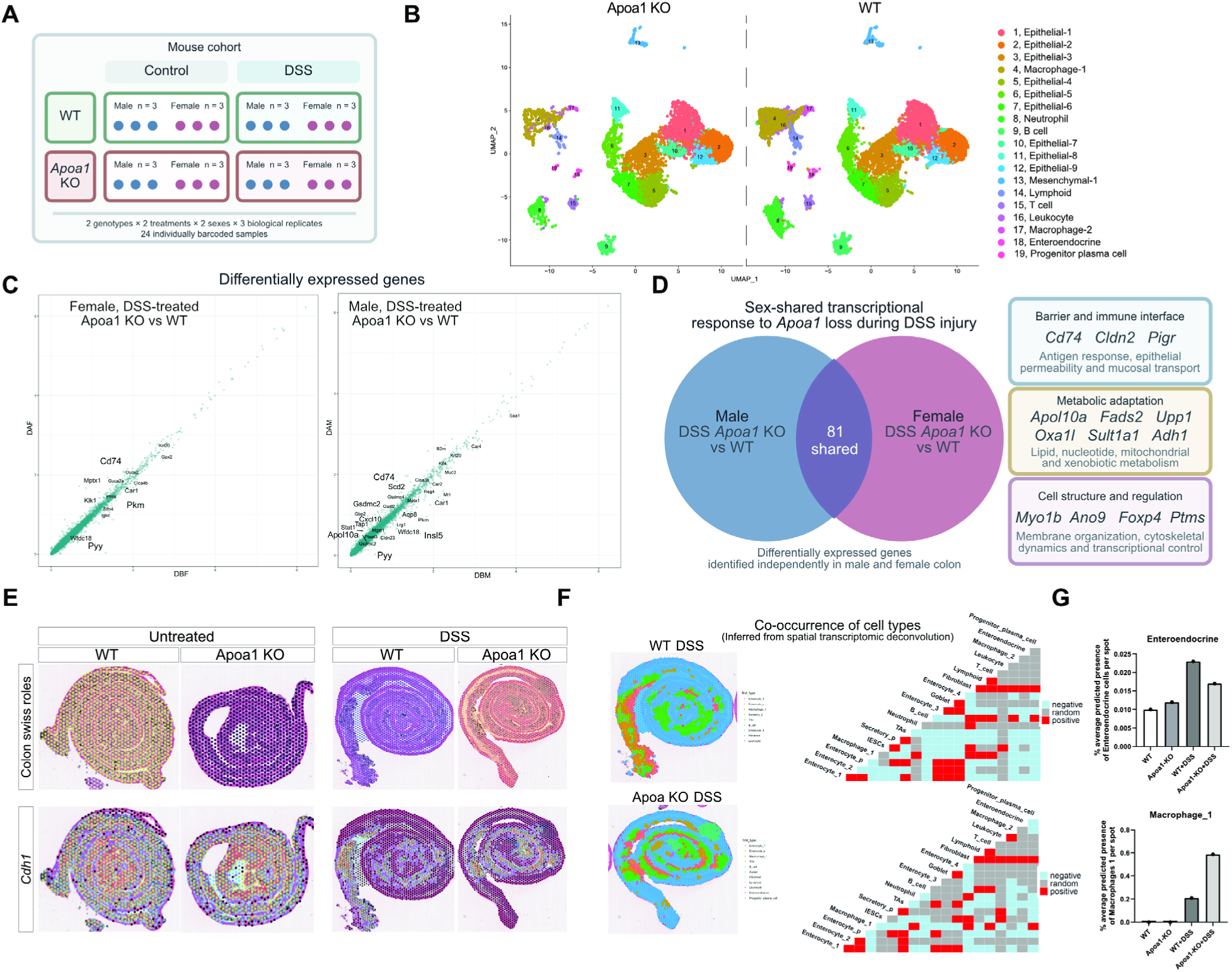
*Apoa1* deficiency is associated with a sex-shared transcriptional response and altered epithelial–immune organization following colonic injury. (A) Experimental design for single-cell RNA sequencing and spatial transcriptomic analysis. Wild-type (WT) and *Apoa1*- knockout mice of both sexes were maintained untreated or exposed to dextran sulfate sodium (DSS). Three biological replicates were included for each sex, genotype and treatment combination, corresponding to 24 individually barcoded samples. (B) Uniform manifold approximation and projection (UMAP) representation of the integrated single-cell RNA-sequencing dataset, displayed separately for *Apoa1*-knockout and WT cells. Colors indicate the annotated epithelial, immune and mesenchymal cell populations listed at right. (C) Gene-level mean-expression comparisons between DSS-treated WT and *Apoa1*-knockout samples, analyzed separately in females and males. Each point represents one gene, and selected differentially expressed genes are labeled. DBF, DSS-treated WT female; DAF, DSS-treated *Apoa1*-knockout female; DBM, DSS-treated WT male; DAM, DSS-treated *Apoa1*-knockout male. (D) Overlap between genes differentially expressed in DSS-treated *Apoa1*-knockout versus WT colon in males and females. Eighty-one genes were identified in both comparisons. Selected shared genes are displayed according to descriptive functional categories encompassing the barrier–immune interface, metabolic adaptation, and cellular structure or regulation. (E) Spatial transcriptomic maps of colon Swiss-roll sections from untreated and DSS-treated WT and *Apoa1*-KO mice. Upper panels show the spatial distribution of transcriptomic clusters; lower panels show *Cdh1* expression, outlining epithelial regions and crypt organization. (F) Spatially predicted cell-type distributions and ISCHIA-based pairwise co-occurrence matrices for DSS-treated WT and Apoa1-KO colon. Positive and negative associations indicate co-occurrence more or less frequent than expected from the individual occurrence frequencies of each cell type; “random” indicates no significant departure from expectation. Spatial analysis units, cell-type presence thresholds and statistical criteria are described in the Methods. (G) Relative predicted spatial representation of enteroendocrine and Macrophage-1 populations in untreated and DSS-treated WT and *Apoa1*-knockout colon. Bars show [mean ± SD/SEM; specify], with individual biological replicates and statistical tests described in the Methods.

To assess the effects of *Apoa1* deficiency in each sex, we compared DSS-treated *Apoa1*-KO and WT samples separately in males and females. These comparisons identified hundreds of differentially expressed genes in each sex (Fig. 4C), with 81 genes shared between the two analyses (Fig. 4D). Shared genes included *Cd74*, *Cldn2* and *Pigr*, linking the response to antigen presentation, epithelial permeability and mucosal immunoglobulin transport. These genes also have established relevance to intestinal injury: CD74 signaling supports epithelial regeneration and mucosal healing, CLDN2 contributes to recovery following colonic damage, and PIGR-dependent secretory immunity influences susceptibility to DSS colitis^20–22^.

The shared gene set additionally included *Apol10a*, *Fads2*, *Upp1*, *Oxa1l*, *Sult1a1* and *Adh1*, encompassing lipid, nucleotide, mitochondrial and xenobiotic metabolism. Other shared genes included *Myo1b*, *Ano9*, *Foxp4* and *Ptms* (Fig. 4D). Thus, the transcriptional consequences of *Apoa1* deficiency extended across barrier-associated, immune and metabolic functions.

Beyond this shared set, the sex-stratified analyses identified stress- and defense-associated genes, including *B2m*, *Cd74* and *Cxcl10*, together with interferon- and cytokine-response genes such as *Stat1*, *Stat2*, *Oasl1*, *Oasl2*, *Gbp2* and *Apobec3*. Differentially expressed genes also included the gasdermin-family members *Gsdmc*, *Gsdmc2* and *Gsdmc3*. These findings further implicated inflammatory and stress-associated transcriptional responses in the injured *Apoa1*-KO colon.

We next used spatial transcriptomics to examine tissue organization following injury. Mapping *Cdh1* expression delineated epithelial regions and crypt organization in untreated and DSS-treated WT and *Apoa1*-KO colon (Fig. 4E). To investigate relationships between epithelial and immune populations, we applied ISCHIA, which identifies cell-type pairs that co-occur more or less frequently than expected from their individual occurrence frequencies^23^. This analysis complements cell-type abundance estimates by assessing preferential spatial association. Applied to our deconvoluted spatial data, using match single cell RNA sequencing datasets (Fig. 4B), it identified co-occurrence patterns involving macrophage-associated and epithelial populations in DSS-treated WT and *Apoa1*-KO tissue (Fig. 4F).

Spatial estimates of cell-type representation showed lower enteroendocrine representation in DSS-treated *Apoa1*-KO tissue than in DSS-treated WT tissue, paralleling the enteroendocrine phenotype observed during development. Conversely, the predicted representation of the Macrophage-1 population, characterized by an inflammation-associated transcriptional profile, was increased in DSS-treated Apoa1-KO colon (Fig. 4G), placing the epithelial changes within an altered inflammatory tissue environment. Together, these analyses associate *Apoa1* deficiency with transcriptional changes detected in both sexes and with altered enteroendocrine and macrophage representation following colonic injury.

#### HOXB7 supports Apoa1 expression in developing colonic tissue

Having characterized the consequences of *Apoa1* deficiency, we next sought to identify transcriptional regulators that could account for the spatially restricted expression of *Apoa1*. We applied SCENIC^24^ to the E18.5 hindgut and DSS-injured adult-colon datasets to infer transcription-factor regulon activity associated with the APOA1-associated epithelial state. This analysis nominated several candidate regulators, including MAF, STAT1, PPARA, NR1H3, MAX, and HOXB7, which we examined by targeted expression analysis (Fig. 5A).

**Figure 5.**
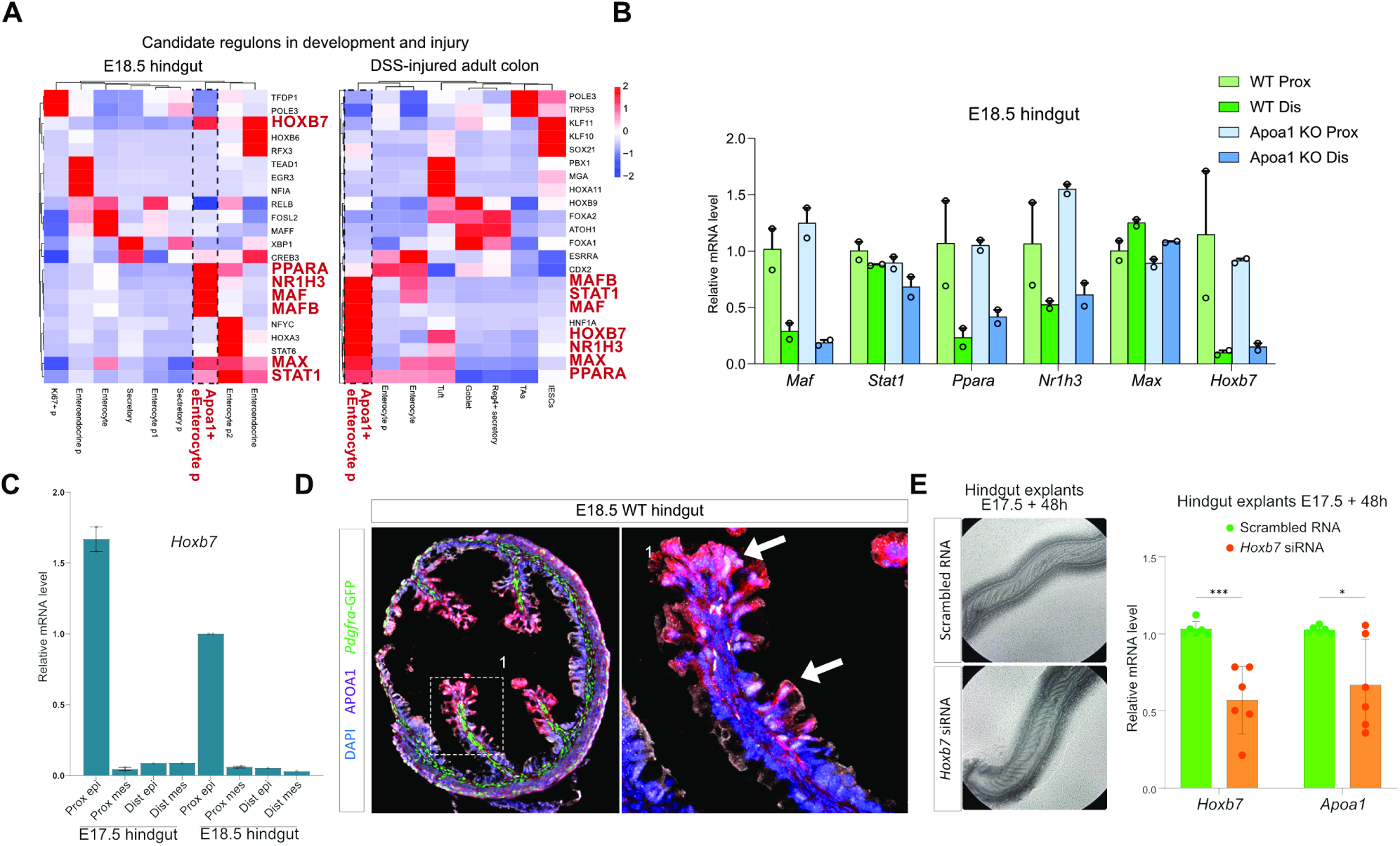
Candidate transcriptional regulators and Hoxb7 perturbation in developing colonic tissue. (A) Heatmaps of SCENIC-inferred regulon activity across annotated cell populations in E18.5 hindgut and DSS-injured adult colon. Selected candidate regulators are highlighted in red. (B) RT-qPCR analysis of *Maf*, *Stat1*, *Ppara*, *Nr1h3*, *Max*, and *Hoxb7* in proximal and distal E18.5 hindgut from wild-type (WT) and Apoa1-knockout (KO) embryos. (C) RT-qPCR analysis of Hoxb7 in sorted epithelial (epi) and mesenchymal (mes) fractions from proximal and distal hindgut at E17.5 and E18.5 using EPCAM antibody. (D) Representative immunofluorescence images of E18.5 WT hindgut showing APOA1 (magenta), HOXB7 (red), Pdgfra-GFP (green), and DAPI (blue). The boxed region is shown at higher magnification on the right. (E) Representative brightfield images and RT-qPCR analysis of E17.5 hindgut explants cultured for 48 h following proximal luminal delivery and electroporation of scrambled control siRNA or Hoxb7-targeting siRNA. Hoxb7 and Apoa1 transcript levels are shown.

RT-qPCR analysis of proximal and distal E18.5 hindgut showed proximal enrichment of several candidates, including *Hoxb7* and *Ppara*. These regional patterns were broadly preserved in both WT and *Apoa1*-KO tissue, indicating that their expression was not dependent on APOA1 (Fig. 5B). We next assessed the tissue compartments in which *Hoxb7* was expressed. Analysis of sorted epithelial and mesenchymal fractions showed that *Hoxb7* expression was concentrated in proximal epithelium, with little expression in distal or mesenchymal fractions. Proximal epithelial *Hoxb7* expression was already evident at E17.5 and was higher than at E18.5 in WT, preceding the prominent developmental increase in *Apoa1* expression (Fig. 5C). Immunofluorescence further localized HOXB7 to the proximal epithelial domain containing APOA1-positive cells (Fig. 5D). Thus, the regional, compartmental, temporal, and spatial correspondence between HOXB7 and APOA1 led us to select HOXB7 for functional testing, while PPARA, MAF, and the other SCENIC candidates remained plausible co-regulators.

To test whether *Apoa1* expression is regulated by HOXB7, we introduced *Hoxb7*-targeting or scrambled control siRNA into the proximal lumen of E17.5 hindgut explants, followed by electroporation and 48 h of culture. *Hoxb7*-targeting siRNA reduced *Hoxb7* transcript levels and was accompanied by lower *Apoa1* expression compared with scrambled controls (Fig. 5E). Thus, reducing *Hoxb7* expression in developing colonic tissue diminished expression of *Apoa1*.

Together, regulon inference, spatial and temporal expression analysis, and functional perturbation support HOXB7 as an upstream contributor to developmental *Apoa1* expression, linking the APOA1-associated epithelial state to a candidate transcriptional regulator shared between development and injury.

## Discussion

Developmental programs reactivated during injury carry potential biological functions as well as molecular signatures of their origin. Here, we identify APOA1 as a functional component of a late fetal colonic epithelial state, connecting its loss to altered mucosal maturation, enteroendocrine organization, and the inflammatory response to adult injury. Building on our identification of developmentally distinct epithelial programs that reappear after damage^7^, these findings advance from recognizing a shared transcriptional identity to testing the contribution of one of its products. They support a broader possibility: programs associated with late developmental maturation may supply functions relevant to tissue organization and differentiation when expressed in the injured adult colon.

This distinction matters for interpreting intestinal plasticity. Studies of extracellular matrix-dependent YAP/TAZ activation and mesenchymal asporin have established mechanisms through which injured epithelium acquires fetal-like properties^1,2^. Development itself, however, comprises successive transitions in epithelial identity, architecture, and function. Assigning an injury-associated program to a defined developmental interval therefore provides a basis for asking what biological activities it might contribute. The late fetal origin of the APOA1-associated state places it in a period of mucosal maturation. Its functional association with developing architecture and endocrine composition suggests that developmental reactivation can be investigated in terms of the tissue-building activities represented within the re-expressed program.

The developmental role of APOA1 extends its established intestinal biology. Earlier work demonstrated apolipoprotein synthesis in human fetal colon and linked regional APOA1 enrichment to resistance to experimental colitis in mice^10,11^. Our findings connect this expression to the maturation of colonic structure. The transient proximal fold phenotype is particularly informative: it is consistent with APOA1 contributing to the pace or efficiency of morphogenesis.

The enteroendocrine phenotype raises the possibility that this environment supports the coordination of tissue growth with lineage maturation. Enteroendocrine identity is shaped by both intrinsic differentiation programs and extracellular signals: temporally resolved lineage analyses have identified successive regulatory transitions, while BMP signaling can alter hormone expression in differentiated enteroendocrine cells^13,25^. Against this background, the relative enrichment of HHEX transcription factor within a smaller NEUROD1-positive compartment suggests that *Apoa1* deficiency affects endocrine populations unevenly. HHEX enrichment in somatostatin-expressing D cells makes subtype balance a relevant direction for further investigation^26^. A local effect on differentiation, survival, or endocrine identity could connect APOA1 availability to this phenotype without requiring APOA1-expressing cells to belong to the endocrine lineage.

WNT5A provides a plausible molecular connection between apolipoprotein biology and the extracellular regulation of epithelial organization. Lipoprotein-mediated transport of biologically active WNT5A in the developing brain established that lipid carriers can participate in morphogen distribution^27^. In the intestine, WNT5A contributes to crypt regeneration through TGF-β signaling and can cooperate with BMP2 in enterocyte differentiation^17,18^. These studies make ligand availability and epithelial responsiveness complementary considerations for interpreting the APOA1–WNT5A relationship. Tissue proximity, altered ligand expression and localization, and the divergent *Fabp1* trajectories in WNT5A-exposed WT and *Apoa1*-KO organoids collectively motivate investigation of both processes. Increased ligand abundance could coexist with altered spatial presentation or a change in the competence of responding cells. APOA1-dependent WNT5A transport is therefore a testable model within a broader relationship that may also involve ligand production, retention, uptake, and epithelial state.

The injury findings place this developmental function within the established protective biology of APOA1. Previous studies demonstrated suppression of intestinal inflammatory signaling by HDL and APOA1, increased colitis susceptibility following *Apoa1* deletion, and reduced monocyte accumulation after treatment with an APOA1 mimetic^11,12,28^. APOA1 mimetics also mitigate intestinal inflammation by limiting the activity and accumulation of proinflammatory lipids, with effects on both macrophages and intestinal epithelial cells^29^. This work provides a mechanistic framework in which altered lipid handling could contribute to the epithelial stress and inflammatory macrophage representation associated with *Apoa1* deficiency in our study. Our analyses bring epithelial state, endocrine representation, and the spatial inflammatory environment into this framework. The transcriptional overlap between male and female comparisons identifies a shared component of the response to deficiency, while the stress- and immune-associated changes suggest that APOA1 availability influences how injured tissue accommodates inflammatory challenge. The recurrence of an enteroendocrine alteration in development and injury is especially informative because it identifies a cellular compartment affected in both settings. Whether this reflects a shared dependence on maturation signals or different consequences of the developmental and inflammatory environments is an important question emerging from the study.

The spatial findings further suggest that the consequences of *Apoa1* deficiency should be considered at the level of tissue organization. Greater representation of inflammation-associated macrophages and their regional association with epithelial populations place the epithelial phenotype within an altered multicellular environment. This creates several plausible routes through which APOA1 could influence recovery: by supporting epithelial differentiation, modifying inflammatory signaling, or changing the conditions that sustain immune-cell accumulation. These processes may reinforce one another. Establishing their temporal relationships will help determine whether epithelial and immune changes represent parallel consequences of APOA1 deficiency or components of a coupled tissue response.

HOXB7 provides a complementary entry point into the regulation of this developmental component. Its anatomical correspondence with *Apoa1* and the explant perturbation results support a positive regulatory contribution during late fetal development. Conceptually, this separates two questions: how a developmental expression domain is established, and what its products contribute once expressed. The HOXB7-associated regulon inferred in injured adult tissue raises the possibility that part of the developmental regulatory machinery is reused during injury. Testing that possibility would distinguish reactivation through shared regulatory inputs from convergence on *Apoa1* expression through different signals. This distinction is central to understanding how closely adult regenerative states reconstruct their developmental counterparts.

The principal mechanistic challenge is to resolve local epithelial function within systemic apolipoprotein biology. Constitutive *Apoa1* deletion markedly reduces circulating HDL, so epithelial production and circulating protein remain potential contributors to the tissue phenotypes^9^. Compartment-specific perturbation and local restoration would distinguish these sources while testing the functional importance of the restricted epithelial domain.

More broadly, this study supports investigating regenerative plasticity through the developmental timing and biological activities of the programs that reappear. A late fetal expression state contains products associated with an epithelium undergoing maturation, and APOA1 provides evidence that at least one such product contributes to tissue architecture and cellular composition. This motivates a framework in which the consequences of developmental reactivation depend on which components are expressed, in which compartments, and at what stage of the injury response. Establishing those relationships would move the field toward a functional understanding of how developmental programs are redeployed in adult tissues.

## Materials and Methods

### Resource availability& lead contact

Further information and requests for resources and reagents should be directed to and will be fulfilled by the lead contact, Hassan Fazilaty

### Mice and experimental design

C57BL/6J wild-type mice were obtained from Charles River Laboratories (strain code 027). Constitutive *Apoa1*-knockout mice (B6.129P2-*Apoa1*^tm1Unc/J; Jackson Laboratory stock no. 002055) were generated as previously described^9^. *Pdgfra*^H2B-eGFP reporter mice were obtained from The Jackson Laboratory (stock no. 007669)^30^. Embryos and postnatal animals were collected at the developmental stages indicated in the figures. Both sexes were included unless otherwise stated. Adult mice were 2-4 month old. All animal experiments were performed in accordance with Swiss regulations and were approved by the Cantonal Veterinary Office of Zurich, Switzerland.

### DSS-induced colonic injury

Acute colonic injury was induced as previously described^7^. Briefly, adult mice received 2.5% dextran sulfate sodium (DSS; MPbio 02160110-CF) in their drinking water for 5 days, whereas control mice received regular drinking water. Body weight and clinical condition were monitored daily. Unless otherwise indicated, colonic tissue was collected 3 days after DSS withdrawal. Distal and medial colon regions were collected for single-cell and spatial transcriptomic analyses. The transcriptomic experiment included untreated and DSS-treated wild-type and *Apoa1*-knockout mice of both sexes, with three biological replicates per sex, genotype and treatment condition performed on different days (24 mice in total).

### Tissue collection and cryosectioning

Embryonic and postnatal hindguts and adult colons were dissected in cold PBS. Adult colons were flushed with PBS to remove luminal contents. For regional analyses, the proximal and distal thirds were collected separately. Embryonic and early postnatal tissues were fixed in 4% paraformaldehyde (PFA) at 4°C for 45–60 min; adult tissues were fixed for 2 h. Following three PBS washes, samples were cryoprotected overnight in 30% sucrose at 4°C. An equal volume of optimal cutting temperature compound (OCT; Tissue-Tek, SA62550-01) was added for 1–4 h before embedding in OCT. Blocks were stored at −70 to −80°C. Sections were cut at 10 µm using a CryoStar NX50 or NX70 cryostat and dried for 1 h at room temperature before staining or frozen storage. Tissue processing for spatial transcriptomics is described separately below.

### Immunofluorescence

Cryosections were washed in PBS and blocked for 1 h at room temperature in PBS containing 5% bovine serum albumin (BSA; Sigma-Aldrich, 10735086001) and 0.2% Tween 20. Primary antibodies were diluted in 3% BSA in PBS and incubated on sections for 1 h at room temperature. Following PBS washes, fluorophore-conjugated secondary antibodies were applied at 1:500 for 1 h at room temperature. Nuclei were counterstained with DAPI, and F-actin was labeled with Alexa Fluor 488 phalloidin where indicated. Sections were washed and mounted in Dako Fluorescence Mounting Medium (Agilent, S302380-2). CHGA staining included antigen retrieval in Tris–EDTA buffer containing 10 mM Tris, 1 mM EDTA and 0.05% Tween 20, pH 9, as described previously^7^. Antibody details are listed in Table 1.

**Table 1.** Antibodies and fluorescent reagents. Entries below reproduce reagents documented in the theses. Application-specific gaps are marked explicitly; they are not completed by substituting antibodies from unrelated experiments.

| Target or reagent | Host or conjugate | Supplier and catalog | Documented use and dilution |
| --- | --- | --- | --- |
| APOA1 | Goat polyclonal | Novus, NB600-609 | IF 1:500 |
| HOXB7 | Rabbit polyclonal | Atlas Antibodies, HPA049940 | IF 1:100 |
| LEF1 | Rabbit monoclonal, C12A5 | Cell Signaling Technology, 2230 | IF 1:100 |
| WNT5A | Rat monoclonal | Bio-Techne, MAB645 | IF 1:100 |
| CHGA | Rabbit polyclonal | Sigma-Aldrich, SAB4200668 | IF 1:500; antigen retrieval |
| E-cadherin | Goat polyclonal | Bio-Techne, AF648 | IF 1:200 |
| EPCAM | Rat monoclonal G8.8, FITC | Invitrogen, 11-5791-82 | FACS and epithelial counterstain, 1:500 |
| EPCAM | Rabbit monoclonal | Abcam, ab213500 | IF 1:200 |
| HHEX | Rabbit monoclonal | Bio-Techne, MAB83771 | IF 1:200 |
| NEUROD1 |  |  | IF 1:200 |
| GCG |  |  | IF 1:200 |
| Anti-goat IgG | Donkey, Alexa Fluor 555 | Invitrogen, A32816 | IF 1:500 |
| Anti-rabbit IgG | Donkey, Alexa Fluor 488 | Invitrogen, A21206 | IF 1:500 |
| Anti-rabbit IgG | Donkey, Alexa Fluor 555 | Invitrogen, A32794 | IF 1:500 |
| Anti-rabbit IgG | Donkey, Alexa Fluor 647 | Invitrogen, A32795 | IF 1:500 |
| Anti-rat IgG | Chicken, Alexa Fluor 647 | Invitrogen, A21472 | IF 1:500; PLA PLUS-probe preparation |
| Phalloidin | Alexa Fluor 488 | Invitrogen, A12379 | 1:1,000 |
| DAPI | Nuclear stain | Invitrogen, D1306 | IF 1:500–1:1,000; sorting 1:1,000 |

### RNAscope RNA in situ hybridization

RNA localization was examined in fixed-frozen cryosections using the RNAscope Multiplex Fluorescent Reagent Kit v2 (Advanced Cell Diagnostics/Bio-Techne), following the fixed-frozen tissue preparation, pretreatment, hybridization and amplification workflow in the manufacturer’s protocol, UM 323100. The documented channel reagents were HRP-C1 (323104) and HRP-C3 (323106), combined with TSA Vivid 570 (323272) and 650 (323273), respectively. Nuclei were counterstained with DAPI, and fluorescence images were acquired by confocal microscopy. The *Apoa1* assay was used to localize transcripts in embryonic hindgut and postnatal colon.

### Whole-mount staining and three-dimensional imaging

Dissected hindguts were opened longitudinally and fixed for 1 h in 4% PFA at 4°C. Samples were blocked for 5 h at 4°C in PBS containing 5% BSA and 1% Triton X-100. Primary antibodies were diluted in PBS containing 3% BSA and 1% Triton X-100 and incubated with tissues for two nights at 4°C. Samples were washed with PBS at 30-min intervals for 12 h and incubated overnight at 4°C with secondary antibodies in the same diluent. Following PBS washes at 30-min intervals for 3 h, tissues were mounted with the luminal surface facing the coverslip in 0.3% low-melting-point agarose in glass-bottom dishes (Thermo Fisher Scientific, 150680), covered with PBS and imaged by confocal microscopy.

### Microscopy and image quantification

Images were acquired using a Leica SP8 inverted confocal microscope at the Center for Microscopy and Image Analysis, University of Zurich. Image processing and measurements were performed in Fiji/ImageJ (2.9.0/1.53t), and three-dimensional reconstructions were generated in Imaris 9.5.0. Image settings were kept comparable for images displayed together. Proximal fold length was measured in tissue sections from the center of the fold base to its luminal tip using the length-measurement tool in Fiji. Whole-mount reconstructions provided complementary visualization of fold architecture.

CHGA-positive epithelial cells were counted manually and reported as cells per section. NEUROD1-positive cells and HHEX/NEUROD1 double-positive cells were evaluated in multiplex images, with GCG staining providing additional endocrine characterization. The HHEX-related measurement was calculated using the numerator and denominator specified in the figure legend.

### APOA1–WNT5A proximity ligation assay

In situ proximity ligation was performed using Duolink reagents (Sigma-Aldrich). An anti-rat PLUS probe was prepared using the Duolink In Situ Probemaker PLUS kit (DUO92009) and chicken anti-rat Alexa Fluor 647 antibody (Invitrogen, A21472). For conjugation, 20 µL antibody at 1 mg/mL was mixed with 2 µL conjugation buffer and incubated overnight at room temperature. Stop reagent (2 µL) was added for 30 min, followed by 24 µL storage solution; conjugated probe was stored at 4°C.

Cryosections were dried for 1 h, post-fixed in 4% PFA for 10 min, permeabilized with 0.2% Triton X-100 in PBS for 15 min, and blocked in Duolink blocking solution for 1 h at 37°C. Goat anti-APOA1 and rat anti-WNT5A were applied for 1 h at room temperature in probe diluent. Following washes in Buffer A, anti-goat MINUS probe (DUO92006) and the prepared anti-rat PLUS probe were applied at 1:10 for 1 h at 37°C. Ligation was performed for 30 min at 37°C with 1× ligation buffer and ligase at 1:40. Amplification was performed for 100 min at 37°C using 1× amplification buffer and polymerase at 1:80 (Duolink In Situ Detection Reagents Red, DUO92008). Sections were washed in Buffer B, counterstained with DAPI and an epithelial marker, and mounted in Dako medium. *Apoa1*-KO tissue served as a biological control, and omission of the ligation step served as a technical negative control.

For quantification, 63× confocal z-stacks were converted to maximum-intensity projections. PLA puncta were counted in Fiji and normalized to the tissue area in each image (Fig. 3G). The epithelial counterstain was used to relate the signal to tissue compartments.

### Isolation of epithelial and nonepithelial fractions

Proximal and distal hindgut regions were dissected separately, minced for approximately 1 min and incubated in 600 µL collagenase D at 2 mg/mL for 1 h at 37°C with agitation at 500 rpm. Cells were pelleted at 500 × g for 3 min, washed in PBS containing 1% BSA and incubated with FITC-conjugated anti-EPCAM (clone G8.8; Invitrogen, 11-5791-82; 1:500) for 30 min on ice. After washing, DAPI was added at 1:1,000 and suspensions were passed through a 40-µm filter. EPCAM-positive and EPCAM-negative fractions were collected using a BD FACSAria III at the Cytometry Facility, University of Zurich. *Epcam* and *Acta2* expression were used to assess compartmental enrichment. RNA was isolated using the mirVana miRNA Isolation Kit (Thermo Fisher Scientific, AM1561). These fractions were used for the regional and developmental expression measurements in Fig. 5C. EPCAM-negative fractions are referred to as nonepithelial; the isolation procedure did not positively select a specific mesenchymal lineage.

### RNA extraction and RT-qPCR

RNA from tissue and explants was extracted using TRIzol (Invitrogen, 15596026), with mechanical disruption by repeated passage through a needle before phase separation. RNA from sorted fractions was extracted using mirVana, and organoid RNA was extracted using the RNeasy Mini Kit (Qiagen, 74104), following the respective manufacturers’ protocols. RNA concentration was measured using a NanoDrop 2000 spectrophotometer. Up to 500 ng RNA was reverse-transcribed in a 10-µL reaction containing 2 µL PrimeScript RT Master Mix (Takara, RR036B), with incubation at 37°C for 15 min followed by 85°C for 5 s.

RT-qPCR was performed in technical duplicate or triplicate on a QuantStudio 3 or 5 instrument. Each 10-µL reaction contained 5 µL PowerUp SYBR Green Master Mix (Thermo Fisher Scientific, A25741), 2 ng cDNA and 0.3 µM each forward and reverse primer. Cycling comprised 50°C for 2 min, 95°C for 5 min, and 50 cycles of 95°C for 15 s and 60°C for 1 min, followed by melt-curve analysis. Relative transcript abundance was calculated from Ct values normalized to *Actb*. Available primer sequences are provided in Table 2.

**Table 2.**
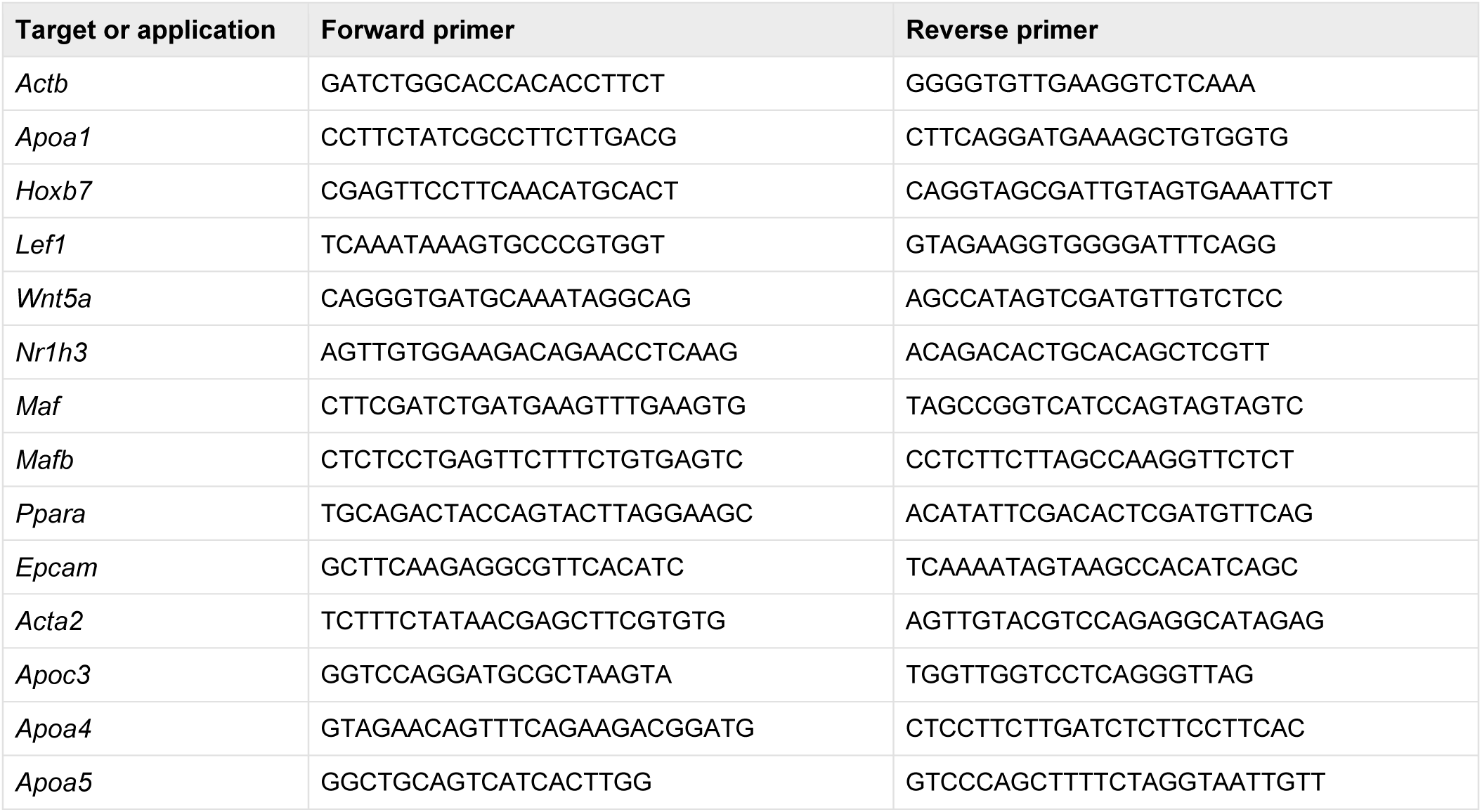
RT-qPCR primers. All sequences are given 5′ to 3′.

### Hindgut explant electroporation

E17.5 WT hindguts were dissected in cold PBS, retaining the cecum to identify orientation, and transferred to cold BGJb medium (Thermo Fisher Scientific, 12591038). The cecum was cut to provide luminal access. Hoxb7-targeting siRNA or MISSION siRNA Universal Negative Control #1 (Sigma-Aldrich, SIC001) was introduced into the proximal lumen at 2 µM in Opti-MEM (Gibco, 11058-021) containing Fast Green (A16520.06). The documented Hoxb7-targeting sequences were 5′-CGAGCCUCUUUCUGUAUAU-3′ and 5′-CCGAAAGACAGAUCAAGAU-3′. An ECM830 Electro Square Porator was used to deliver five unipolar 47-V pulses of 50 ms at 1-s intervals. The proximal tissue was electroporated in two orientations, rotating the electrode arrangement by 180° between applications.

Explants were cultured for 48 h at 37°C on Transwell polycarbonate inserts in six-well plates (Corning, CLS3428), with 700 µL medium beneath the insert and 300 µL above. BGJb medium was supplemented with ascorbic acid, using the previously described^7^ concentration of 0.1 mg/mL. Whole explants were collected for RNA extraction, and *Hoxb7* and *Apoa1* expression was measured by RT-qPCR. The source analysis assessed knockdown efficiency first and evaluated *Apoa1* in the subset exhibiting reduced *Hoxb7* expression.

### Colonic organoid culture and WNT5A treatment

Organoids were established from adult female WT and *Apoa1*-KO colons. Dissected colons were washed, opened longitudinally, cut into small pieces and washed repeatedly with PBS. Tissue fragments were incubated in Gentle Cell Dissociation Reagent (STEMCELL Technologies, 100-0485) for 30 min at room temperature with rocking. Crypt-containing fractions were released by pipetting in PBS containing 0.2% BSA and passed through a 70-µm filter. Fractions containing intact crypts with limited debris were selected, centrifuged at 290 × g for 5 min at 4°C, washed and embedded in Matrigel (Corning, 356255) in prewarmed 24-well plates. Matrigel was polymerized for 10 min at 37°C before medium addition.

Complete medium comprised Advanced DMEM/F12 (12634-010), 1× GlutaMAX (35050061), 10 mM HEPES (15630080), 1× N2 (17502-048), 1× B27 (17504-044), 1 mM N-acetyl-L-cysteine (Sigma-Aldrich, A9165), 50 ng/mL EGF (PMG8041), 1 µg/mL R-spondin 1 (PeproTech, 315-32) and 0.5 nM Wnt surrogate (N001). Cultures were maintained at 37°C with 5% CO2 and passaged mechanically approximately every 3 days. Following fragmentation and washing, organoids were re-embedded in 40-µL Matrigel domes.

For WNT5A treatment, two domes were plated per well and cultured for 24 h in complete medium. Medium was then replaced with medium lacking R-spondin 1 and Wnt surrogate and supplemented with recombinant human/mouse WNT5A (Bio-Techne, 645-WN-010/CF). Organoids were collected after 12, 24, 48 or 72 h. Untreated control cultures were collected at 12 h. RNA was isolated using RNeasy, and *Fabp1* expression was measured by RT-qPCR (Fig. 3I); *Lef1* was measured in the same experimental series.

### Single-cell sample preparation and MULTI-seq

Single-cell RNA sequencing was performed on E18.5 WT and *Apoa1*-KO hindgut and on the 24 adult samples described above. Embryonic and adult samples were barcoded using MULTI-seq^31^ (Sigma-Aldrich, LMO001) to retain sample identity after pooling. Each sample received a distinct oligonucleotide barcode associated with a membrane anchor and co-anchor; excess reagents were removed before pooling. Gene-expression and sample-barcode libraries were prepared separately for sequencing.

### Single-cell library preparation and computational analysis

Barcoded suspensions were processed using the 10x Genomics Chromium platform, and gene-expression libraries were sequenced together with their corresponding MULTI-seq barcode libraries. Gene-expression reads were aligned to the mouse reference transcriptome and converted to gene-by-cell count matrices. Sample-barcode counts were used to assign cell identities to their original biological samples. Expression matrices were filtered, normalized and integrated for visualization, and principal-component analysis and UMAP were used to represent the data. Clusters were annotated using lineage-marker expression.

Gene-expression comparisons were performed between WT and *Apoa1*-KO E18.5 epithelium and separately between DSS-treated WT and *Apoa1*-KO samples within each sex. In the endocrine-marker dot plot, dot size represents the fraction of cells expressing each gene and color represents the plotted mean-expression scale. The male and female DSS differential-expression lists were intersected by gene identity, yielding the 81-gene overlap. Selected genes were grouped into descriptive functional categories for display.

### SCENIC regulon analysis

Candidate regulatory programs in E18.5 hindgut and DSS-injured adult colon^7^ were examined using SCENIC^24^. The workflow infers transcription-factor-associated coexpression modules, refines candidate target sets using cis-regulatory motif enrichment, and scores regulon activity in individual cells using AUCell. Regulon profiles were summarized across the annotated epithelial populations to identify candidates associated with the APOA1-marked state.

### Spatial transcriptomics

Spatial gene-expression profiling was performed on colon Swiss-roll sections from untreated and DSS-treated WT and *Apoa1*-KO mice using the 10x Genomics Visium HD platform. Spatially indexed expression data were registered to the tissue images and used to visualize epithelial organization, including the distribution of *Cdh1* expression. Cell-type composition was estimated using annotated single-cell expression profiles as the reference. The resulting spatial cell-type maps were used for representation and co-occurrence analyses.

### Spatial co-occurrence analysis

Cell-type co-occurrence was analyzed using ISCHIA (Identifying Spatial Co-occurrence in Healthy and InflAmed tissues)^23^. Estimated cell-type abundances were converted to presence–absence calls at the analyzed spatial resolution. Pairwise observed co-occurrence was compared with the expectation based on the marginal occurrence frequencies of each population. Associations were classified as positive, negative or indistinguishable from the model expectation and displayed as pairwise matrices. This analysis assesses preferential sharing of spatial locations while accounting for population prevalence. Analyses were interpreted at the spatial-bin or spot resolution used for deconvolution.

### Statistical analysis and data availability

Non-transcriptomic analyses were performed in GraphPad Prism (versions 9 and 10). Proximal fold lengths were compared between WT and *Apoa1*-KO groups at each developmental stage using unpaired t-tests. Statistical tests, sample numbers and measures of central tendency and dispersion are specified in the corresponding figure legends. Technical qPCR wells, image fields and sections are distinguished from independent biological samples.

## Acknowledgments

We sincerely thank Konrad Basler for immense support including financial support for this work, George Hausmann, Erich Brunner, and Luciano Rago for treasured discussions, and members of the Basler lab for valuable input. We thank Susanna Salas for administrative assistance, Werner Wolz for IT support and Tosca Dalessi for technical assistance. We also acknowledge the technical support from the Functional Genomics Center Zurich, especially Catharine Aquino. We also thank the Office for Animal Welfare and 3R of the University of Zurich, mainly Michaela Thallmair, Corina Berset, Nicole Wildner and Paulin Jirkof for supporting animal experimentation procedures. This work was supported by: Swiss National Science Foundation (192475). T.V. was supported by Czech Science Foundation grant 25-17207S. H.F. was supported by Swiss National Science Foundation grant (CRSK-3_237773), Enrst Hadorn Transitional fellowship, University of Zurich FAN fellowship, the University of Zurich GRC career grant (2024_Q1_CG_001), the Forschungskredit of the University of Zürich (FK-21-119) and a grant from the University and Medical Faculty of Zürich and the Comprehensive Cancer Center Zürich.

## Author contributions

H.F. conceived and supervised the study, acquired funding, performed experiments, curated and analyzed the data, coordinated the project, validated the findings, contributed to data visualization, and wrote the manuscript. A.B., V.G.K.V. and D.K. contributed equally to experimental work and participated in methodology development and data visualization. T.V., G.M., C.B. and M.D.B. contributed to experiments, methodology development and data analysis. P.D. contributed to experiments and data visualization. A.M. provided resources.

